# Slow Dissociation of Nitazenes from the *μ*-Opioid Receptor Underlies the Challenge of Overdose Reversal

**DOI:** 10.64898/2026.04.14.718203

**Authors:** Joseph Clayton, Laura B. Kozell, Amy J. Eshleman, Shelley H. Bloom, William E. Schutzer, Atheir I. Abbas, Lidiya Stavitskaya, Jana Shen

## Abstract

Nitazenes are driving a wave of overdose deaths in the United States and Europe and often require additional doses of naloxone to reverse. To understand the molecular basis, we conducted a joint experimental and simulation study of three common nitazenes, eto-, etodes-, and protonitazene. Radioligand experiments demonstrated that all three nitazenes display higher receptor affinity and longer dissociation half-lives than fentanyl. Notably, protonitazene dissociates slower than carfentanil and its displacement requires fourfold higher antagonist concentrations. The observed trend in nitazene half-lives is recapitulated by molecular dynamics simulations, which suggest that kinetics is controlled by specific interactions with two receptor subpockets. A newly published cryo-EM structure of fluetonitazene-*μ*OR complex confirms the predicted interactions, including a *π*-hole bond between the nitro group and Tyr^1.39^, a residue recently shown to modulate *μ*OR signaling bias. Our findings suggest slow receptor dissociation as a key factor challenging overdose reversal. The mechanistic insights have implications for understanding opioid toxicity and designing more effective countermeasures.

---

Synthetic opioids continue to drive drug overdose deaths in the US. Since 2019, a class of new synthetic opioids known as the nitazenes has caused a rising number of deaths in the US^[1]^ and Europe^[2]^. Nitazenes contain a benzimidazole core substituted by an ethylamine and a benzyl group (Figure 1a). According to the US Drug Enforcement Administration’s March 2026 Orange Book, 21 nitazenes have been placed under Schedule I controlled substances, including proto-, eto-, and etodesnitazene^[3]^. Several experimental studies demonstrated that some nitazene derivatives (e.g., etonitazene) are more potent than fentanyl *in vitro* ^[4–8]^ and *in vivo*. ^[5]^ A 2023 analysis of an emergency department cohort found that overdosed patients testing positive for nitazenes received more doses of naloxone compared to cases involving fentanyl alone.^[9]^ Naloxone is a commonly used opioid reversal agent approved by the US Food and Drug Administration (FDA). Three more recent studies also showed that half of the admitted patients testing positive for nitazenes required additional doses of naloxone.^[10–12]^ Since opioids (agonists) compete with naloxone (antagonist) for binding to the *μ*-opioid receptor (*μ*OR), we hypothesized that some nitazenes exhibit longer residence time than fentanyl, thereby requiring a higher dose of naloxone for reversal.

**Figure 1.**
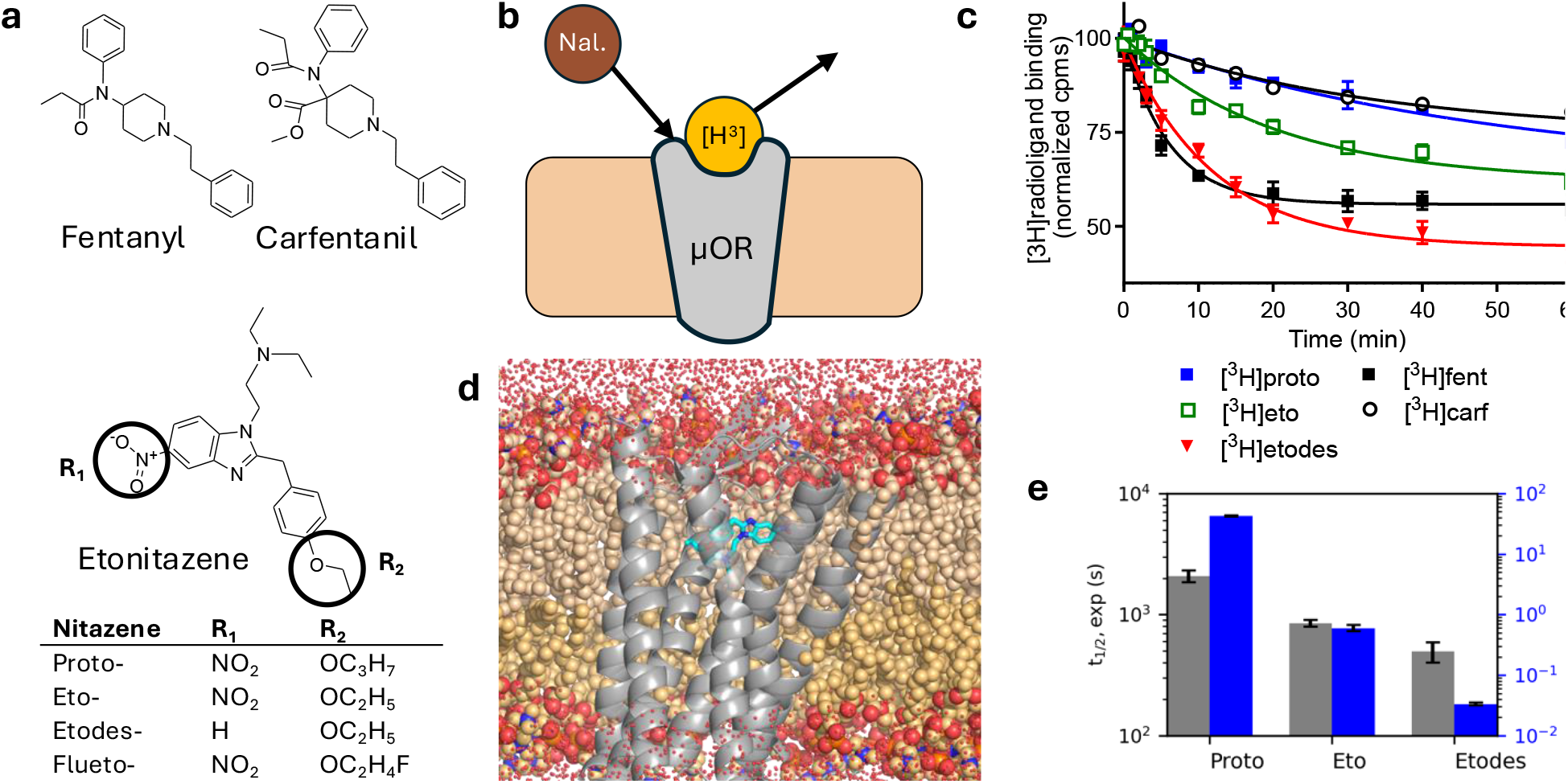
Overview of the experimental and computational investigation of three common nitazenes. **a)** Structures of two fentanyls and three nitazenes studied here. Fluetonitazne (shown in Figure 2d) is also listed. **b)** Schematic of the radioligand dissociation assay. The fraction of radiolabeled ligand (yellow, e.g., a nitazene) remaining bound to the receptor is monitored over time in the presence of unlabeled naloxone (brown). **c)** Time course of the measured counts per minute (cpm, normalized to the initial value) for the [^3^H]-labeled ligand bound to *μ*OR in the presence of 10 nM naloxone. Data for proto-, eto-, etodesnitazene, fentanyl, and carfentanil are shown. Curve represents the best fit to the one phase exponential decay model, from which the half-life (*t*_1*/*2_) is derived (Table 1). **d)** Visualization of the simulation system of protonitazene (cyan) bound to *μ*OR. **e)** Experimental *t*_1*/*2_ values (gray) in comparison to simulation estimates (blue) for the three nitazenes.

To test the above hypothesis, we employed experimental and molecular dynamics (MD) approaches to investigate three common nitazenes: proto-, eto-, and etodesnitazene, which differ by the frequently modified nitro and alkoxy substitutions (Figure 1a). We determined the affinities, kinetics, and naloxone/nalmefene inhibitory constants for the nitazenes and two reference opioids, fentanyl and carfentanil. All three nitazenes displayed higher affinities, dissociation half-lives, and higher naloxone/nalmefene *K*_*i*_ values compared to fentanyl. MD simulations recapitulated the observed trend in dissociation kinetics and identified specific nitazene-receptor interactions, in striking agreement with the newly published cryo-EM structure of fluetonitazene-*μ*OR complex.^[13]^ Intriguingly, a *π*-hole bond (absent in other opioids) is formed with Tyr^1.39^, which has been recently found to modulate *μ*OR signaling bias. ^[14]^

**Table 1.** Receptor binding affinity and kinetics of selected nitazenes and reference agonists and antagonists^*a*^.

| Compound | $K_d$<br>(nM) | $k_{on}$<br>($\text{nM}^{-1}\text{min}^{-1}$) | $t_{1/2}$<br>(min) | Cal. $t_{1/2}$<br>(sec) | Naloxone $K_i$<br>(nM) | Nalmefene $K_i$<br>(nM) |
| --- | --- | --- | --- | --- | --- | --- |
| Protonitazene | $0.164 \pm 0.060$ | $0.93 \pm 0.33$ | $34.5 \pm 2.7$ | $18.3 \pm 0.3$ | $16.0 \pm 3.9$ | $1.93 \pm 0.25$ |
| Etonitazene | $0.082 \pm 0.002$ | $1.12 \pm 0.13$ | $14.2 \pm 0.6$ | $0.26 \pm 0.02$ | $11.8 \pm 2.5$ | $1.70 \pm 0.24$ |
| Etodesnitazene | $0.473 \pm 0.047$ | $0.69 \pm 0.14$ | $8.3 \pm 1.1$ | $0.0144 \pm 0.0006$ | $4.2 \pm 1.2$ | $1.12 \pm 0.32$ |
| Fentanyl | $0.602 \pm 0.063$ | $0.68 \pm 0.14$ | $4.27 \pm 0.53$ | | $2.1 \pm 0.2$ | $0.27 \pm 0.07$ |
| Carfentanil | $0.049 \pm 0.004$ | $0.67 \pm 0.04$ | $22.9 \pm 1.8$ | | $4.7 \pm 0.7$ | $0.49 \pm 0.05$ |
| Naloxone | $0.711 \pm 0.077$ | $3.96 \pm 0.67$ | | | | |
| Nalmefene | $1.490 \pm 0.790$ | $2.43 \pm 0.30$ | | | | |

<sup>a</sup> $K_d$ , $k_{on}$ , and $t_{1/2}$ were from saturation binding, association, and dissociation experiments using radioligands. Saturation and association experiments were performed in the presence of 1 $\mu\text{M}$ unlabeled naloxone, while dissociation experiments were performed in the presence of 10 nM unlabeled naloxone. Calculated $t_{1/2}$ was based on Poisson fitting using the dissociation times from 15 metadynamics trajectories (Table S2). The $K_i$ of unlabeled naloxone or nalmefene was determined by competitive displacement using radioligand concentrations at or above the $K_d$ value for each ligand.

Using radioligand binding assays at the human *μ*OR, the affinities of proto-, eto-, and etodesnitazenes were determined and bench-marked against the well-studied ultra-potent synthetic opioids fentanyl and carfentanil as well as the opioid reversal agents naloxone and nalmefene (Figure 1a and Table 1, Figure S1). Details are given in Supplementary Methods. The *K*_d_ values follow the order: nalmefene (1.49 nM) > naloxone (0.71 nM) > fentanyl (0.60 nM) > etodesnitazene (0.47 nM) > protonitazene (0.16 nM) > etonitazene (0.082 nM) > carfentanil (0.049 nM), demonstrating that all three nitazenes bind the receptor more strongly than fentanyl, with etodes-being the weakest and etonitazene the strongest. This affinity trend corroborates the recent *K*_i_ data based on displacement of [^3^H]DAMGO binding:^[6]^ fentanyl (1.26 nM) > etodesnitazene (1.02 nM) > protonitazene (0.30 nM) > etonitazene (0.21 nM). The binding affinity trend of etonitazene > protonitazene > fentanyl also matches the *in vitro* functional potency trend (EC_50_)^[7]^ based on cAMP inhibition and *β*-arrestin2 recruitment. It is worth noting that nalmefene displays twofold weaker binding affinity than naloxone, which has a slightly weaker affinity than fentanyl.

Next, we determined the kinetic constants of the nitazenes and reference compounds using radioligand association and dissociation assays (Figure 1b-c and Table 1, Figure S2-S3). The obtained *k*_on_ values of the three nitazenes are comparable to each other (within measurement error) and similar to those of fentanyl and carfentanil. However, the antagonists naloxone and nalmefene bind the receptor significantly faster than the nitazenes and fentanyls, with the *k*_on_ values of 3.96 and 2.43 nM^-1^min^-1^, respectively.

Using dissociation binding assays, the half-lives (*t*_1/2_) of receptor dissociation for proto-, eto-, and etodesnitazenes were determined as 34.5, 14.2, and 8.3 min, respectively (Figure 1c and Table 1). Notably, all three values exceed that of fentanyl (4.3 min); importantly, protonitazene’s *t*_1/2_ also surpasses that of carfentanil (28.6 min). These measurements were conducted in the presence of 10 nM unlabeled naloxone, approximating the plasma concentration of 18 nM achieved with the standard 4 mg intranasal reversal dose.^[15]^ To confirm the rank order of half-lives among the nitazenes, additional experiments were performed at higher naloxone concentrations (Table S1). As expected, at a higher naloxone concentration of 100 nM, the *t*_1*/*2_ values of all three nitazenes slightly decrease; however, the rank order remains, with protonitazene’s *t*_1*/*2_ (25.9 min) being more than twice that of etonitazene and more than four times that of etodesnitazene. Increasing the naloxone concentration further to 1 *μ*M produced no meaningful changes in the *t*_1*/*2_values, within measurement errors. To confirm that protonitazene dissociates slower than fentanyls, measurements were conducted with 10 *μ*M. Notably, even at this physiologically irrelevant naloxone concentration, protonitazene’s *t*_1*/*2_ (23.5 min) remains more than six times that of fentanyl (3.7 min) and slightly longer than carfentanil (19.7 min). These data demonstrate that all three nitazenes dissociate more slowly from the receptor than fentanyl and protonitazene dissociates even slower than carfentanil.

To further probe nitazene’s extended interactions with the receptor in comparison to fentanyls, we determined the *K*_*i*_ of naloxone and nalmefene by competitive displacement of a nitazene or a fentanyl at the *μ*OR (Table 1, Figure S4-S5). The naloxone and nalmefene *K*_*i*_ ‘s for proto-, eto-, and etodesnitazenes follow the same order as their dissociation *t*_1*/*2_ values and are all significantly higher than the corresponding *K*_*i*_ for fentanyl. Importantly, the naloxone *K*_*i*_ for proto- or etonitazene is respectively 3.4 or 2.5 fold higher than that for carfentanil, while the nalmefene *K*_*i*_ for all three nitazenes, including etodesnitazene, is 2.3–3.9 fold higher than that for carfentanil.

The above *K*_*i*_ data demonstrate that a higher concentration of naloxone or nalmefene is required for *in vitro* displacement of proto-, eto-, or etodesnitazene from the receptor compared to fentanyl. Strikingly, displacement of protonitazene requires a higher concentration of naloxone or nalmefene than carfentanil, which is the most potent commercially available opioid. Carfentanil overdose cases required additional doses of naloxone for resuscitation.^[16]^ It is also noteworthy that whereas naloxone displaced etodesnitazene and carfentanil at comparable concentrations, a higher concentration of nalmefene was required to displace etodesnitazene.

Recently, we investigated the putative binding poses of a series of nitro-containing and nitro-less nitazenes at *μ*OR through a comprehensive analysis of ligand-bound crystal or cryo-EM structures of *μ*OR, multi-template consensus docking, and advanced MD simulations.^[17]^ We found that *μ*OR’s central cavity contains three subpockets (SPs) capable of recognizing a ligand: SP1 (transmembrane helix or TM2 and TM3), SP2 (TM1, TM2, and TM7), and SP3 (TM3 and TM5). Similar to the indole group of mitragynine pseudoindoxyl (PDB 7T2G),^[18]^ nitazene’s benzimidazole moiety (with or without a nitro substitution) forms specific interactions with residues in SP2 (Figure 2a,b), while the alkoxy tail interacts with residues in SP3 (Figure 2a,c). In contrast, fentanyl’s phenethyl group interacts with SP1 and its propanamide group interacts with SP3 (PDB 8EF5).^[19]^

**Figure 2.**
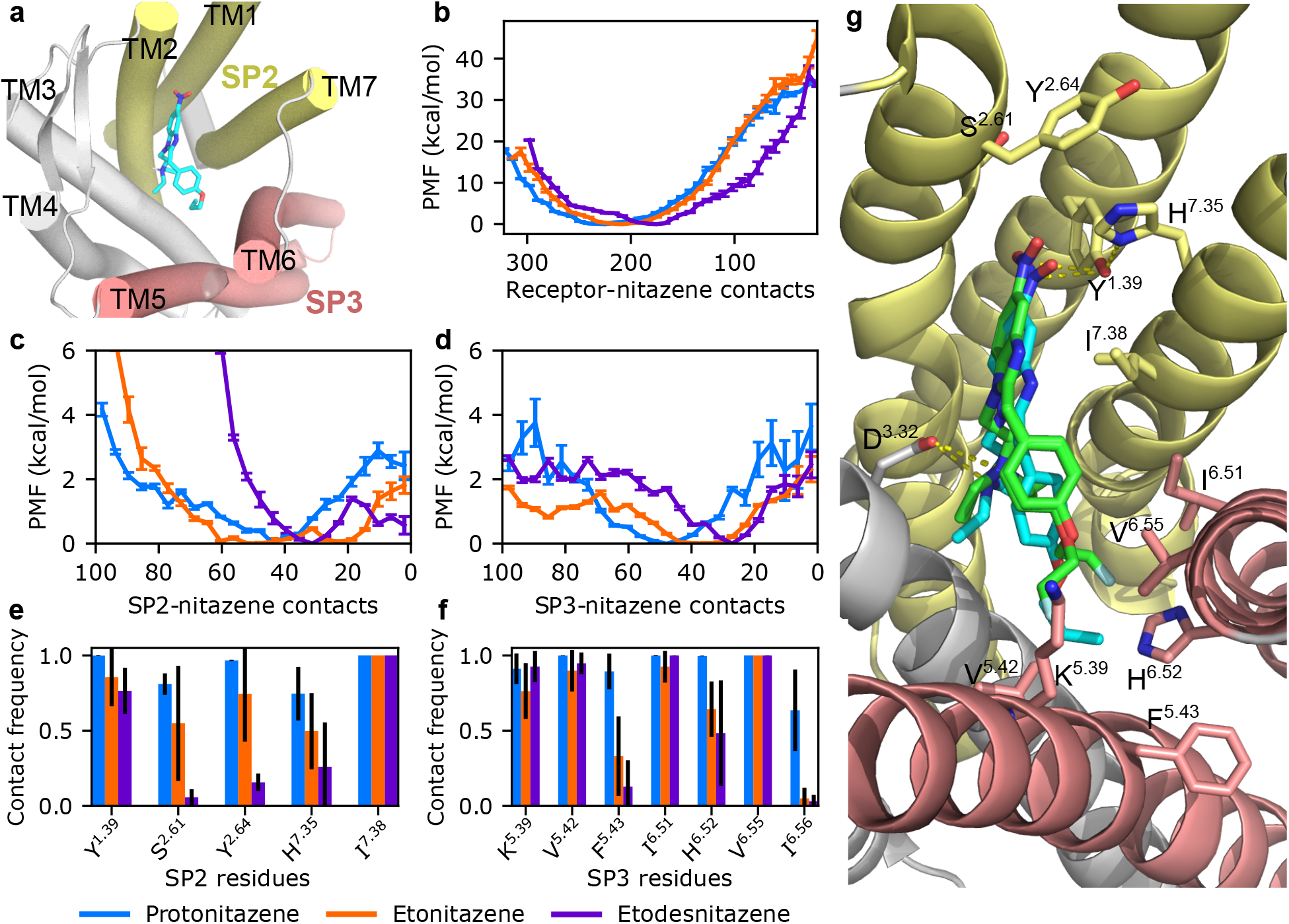
*μ*OR–nitazene interactions are modulated by the nitro and alkoxy substitutions. **a.** Visualization of protonitazene (cyan) within *μ*OR, interacting with subpockets SP2 (TM1, TM2, TM7 in yellow) and SP3 (TM5, TM6 in coral). **b, c, d**. Potential mean force (PMF, also known as free energy profile) as a function of the total number of atomic contacts between *μ*OR (**b**), SP2 (**c**), or SP3 (**d**) and a nitazene. Data for proto-, eto-, and etodesnitazene are shown in blue, orange, and purple, respectively. Standard error are shown. Contact calculations and reweighting protocol are given in Simulation Methods. **e, f**. Contact frequency between residues in SP2 (**e**) or SP3 (**f**) and a nitazene. The final 100 ns of each 500 ns cMD simulations were combined. Only residues with ≥50% contact frequency for at least one nitazene are shown. Residues are numbered according to the Ballesteros-Weinstein scheme. ^[26]^ **g**. A zoomed-in view of simulated protonitazene (cyan) pose overlaid on the cryo-EM structure of fluetonitazene (green) bound *μ*OR (PDB ID: 9O36).^[13]^ Two conformations that mainly differ in the fluorinated ethoxy tail are displayed. The simulated protonitazene pose shown is the representative structure obtained by clustering the final 100 ns of trajectory frames using a ligand RMSD threshold of 2.0 Å, after alignment to the receptor backbone atoms. The *π*-hole bond with Tyr^1.39^, the salt-bridge with Asp^3.32^, and the hydrogen-bond between Tyr^1.39^ and His^7.35^ are indicated by dashed lines. Protonitazene and fluetonitazene interact with the same set of SP2 and SP3 residues (labeled and shown in stick models).

To investigate the mechanistic basis for the extended residence time of protonitazene relative to eto- and etodesnitazene, we performed well-tempered metadynamics simulations^[20,21]^ using a protocol that has been validated for fentanyl and derivatives.^[22,23]^ Following 300 ns conventional molecular dynamics (cMD) simulations to relax the docked structure, 15 metadynamics trajectories were launched to estimate the dissociation half-life. We employed OpenMM (v8.0)^[24]^ package, with the PLUMED (v2.7) plug-in^[25]^ for controlling metadynamics. A time-dependent bias potential was deposited along two collective variables: the ligand’s *z* position relative to that of the receptor’s orthosteric binding pocket and the receptor-ligand coordination number.^[22,23]^ Details are given in Supplementary Methods.

The calculated dissociation *t*_1*/*2_ values of proto-, eto-, and etodesnitazene (18.3, 0.26, and 0.014 s, respectively) match the experimental trend: proto-> eto-> etodesnitazene, although the absolute values are two orders of magnitude smaller than the experimental estimates (Table 1 and Figure 1e, Table S2). This discrepancy may be attributed in part to the missing N-terminal tail in the *μ*OR structure (PDB ID: 7T2G)^[18]^ known to interact with the ligand and in part to the difference between the simulated microscopic dissociation events and macroscopic measurement.

We hypothesized that the dissociation half-life is correlated with receptor-ligand interactions. To test it, we first calculated the free energy as a function of the total number of receptor-ligand atomic contacts defined as receptor-ligand heavy atom pair with a distance not exceeding 4.5 Å. Frames from the metadynamics dissociation trajectories were reweighted. The resulting free energy profiles (also known as potential of mean force) for proto-, eto-, and etodesnitazene display minima at contact numbers of 224, 208, and 176, respectively (Figure 2b), demonstrating that protonitazene forms the largest and etodesnitazene the smallest number of stable interactions, which is consistent with the descending order of their dissociation half-lives.

Next, we examined nitazene-receptor interactions within the two SPs. The free energy profile along the number of atomic contacts between each nitazene and SP2 or SP3 was calculated (Figure 2c and 2d). We hypothesized that the presence of the nitro group (proto- and etonitazene) allows for interactions within SP2 that are not available for nitro-less etodesnitazene. Indeed, the free energy pro-files demonstrate that eto- and protonitazene samples stable states with 40–60 SP2 contacts, whereas these states incur a free energy penalty up to 6 kcal/mol for etodesnitazene (Figure 2c). Notably, although proto- and etonitazene share the same nitrobenzimidazole group, the latter appears more flexible within SP2, sampling states with contact numbers below 40, reaching as low as 20.

We further hypothesized that an additional methylene group in the alkoxy tail (i.e., proto-vs. etonitazene allows for more interactions within SP3. Indeed, the free energy profile along the number of contacts within SP3 (Figure 2d) shows a minimum at 48 for protonitazene and a broader minimum spanning 30–42 for etonitazene while etodesnitazene shows a minimum at 27. Therefore, both SP2 and SP3 interactions modulate the binding kinetics of nitazenes. Protonitazene forming the most stable contacts with both subpockets may explain why it has the longest *t*_1*/*2_ compared to eto- and etodesnitazene. The free energy profiles (Figure 2c and 2d) also suggest some degree of cooperativity between the interactions at SP2 and SP3 – more stable interactions at SP3 (e.g., for protonitazene) may stabilize interactions at SP2.

To corroborate the metadynamics-based analysis and test the hypothesis of cooperativity between the two subpockets, we examined residue-specific interactions from three independent 500-ns equilibrium cMD simulations for each nitazene (Figure S6). Nitazeneresidue interactions, defined as those with contact frequencies of at least 50% in the combined trajectories by at least one nitazene, were analyzed. While all three nitazenes engage the same set of residues within subpockets SP2 and SP3, protonitazene displays the highest frequencies and greatest stabilities for its interactions within SP2 and SP3 relative to eto- and etodesnitazene (Figure 2e and 2f), consistent with the metadynamics-based contact analysis. Within SP2, all three nitazenes form persistent van der Waals contacts with Ile^7.38^. However, the interactions with Ser^2.61^, Tyr^2.64^, and His^7.35^ are most stable for protonitazene, less stable for etonitazene, and nearly abolished for etodesnitazene (occupancy below 25%, Figure 2e, Figure S7). Although protonitazene and etonitazene share the same nitrobenzimidazole at SP2, the more stable interactions with protonitazene suggest positive allosteric coupling between the polar SP2 and hydrophobic SP3. The most intriguing interaction at SP2 is between the electron-deficient nitro nitrogen and the electron-rich hydroxyl oxygen of Tyr^1.39^ (Figure 2g), which represents a *π*-hole bond.^[27,28]^ First proposed in our docking based modeling work for nitazenes,^[17]^ this *π*-hole bond is more stable for protonitazene than etonitazene during simulations (Figure 2e). Interestingly, while Tyr^1.39^ is involved in the *π*-hole bond, it also forms a hydrogen-bond with His^7.35^ (Figure 2g). Note, etodesnitazene also engages Tyr^1.39^ but through van der Waals contacts.

Within SP3, all three nitazene form persistent van der Waals interactions with Val^6.55^, while interactions with Lys^5.39^, Val^5.42^ and Ile^6.51^ are slightly less stable for eto- and etodesnitazene (Figure 2f, Figure S8). Due to the extra methylene group in protonitazene, the propyl group reaches to the back of the SP3, forming hydrophobic interactions with Phe^5.43^ and His^6.52^, and occasionally with Ile^6.56^ (Figure S5 and S6). As expected, the interactions with His^6.52^ and Phe^5.43^ are significantly weaker for eto- and etodesnitazene, which rarely interact with Ile^6.56^.

Shortly before this work was submitted, a cryo-EM structure model of fluetonitazene bound *μ*OR was published^[13]^ (PDB: 9O36, Figure 2g). Strikingly, the cryo-EM model displays a *π*-hole interaction between the nitro group and Tyr^1.39^, with a nitrogen-to-hydroxyl oxygen distance of 3.0 Å. Furthermore, the representative binding pose of protonitazene from the simulations closely resembles that of fluetonitazene in the cryo-EM structure, interacting with an identical set of SP2 and SP3 residues (Figure 2e and 2f with the exception of Ile^6.56^ which transiently interact with protonitazene in the simulations).

In summary, this work used radioligand binding experiments and molecular dynamics simulations to investigate three common nitazenes. Compared to fentanyl, all three nitazenes display higher receptor affinity, longer dissociation half-life, and their displacement requires a higher concentration of naloxone or nalmefene. Most notably, protonitazene dissociates more slowly than carfentanil and displacement requires four times the concentration of naloxone or nalmefene. We found that the naloxone or nalmefene concentrations needed for opioid displacement are correlated with the half-lives, rather than the receptor affinities. These data provide a molecular rationale for recent clinical observations that patients testing positive for nitazenes often required additional doses of naloxone for effective reversal.^[9–12]^

The MD simulations reproduced the experimental trend in dissociation kinetics and suggest that nitazene’s half-life is modulated by the nitro–SP2 and alkoxy–SP3 interactions, which explains why the half-life of protonitazene is longer than eto- or etodesnitazene. Given that proto- and etonitazene share the nitrobenzimidazole motif, but differ in the alkoxy tail length, the reduced interactions of etonitazene within both subpockets suggests a cooperative allosteric effect. Although the trend in the simulation-estimated half-lives is consistent with experiment, the absolute values are 2–4 orders of magnitude lower. We attribute this discrepancy to the absence of the N-terminal loop in the *μ*OR structure used, as well as to the substantial timescale mismatch between simulations on the order of 100 ns per trajectory and real-world dissociation events occurring on the order of 100–1000 s. We note that a comparison of the current nitazene simulations and our previous metadynamics dissociation simulations of fentanyl and its derivatives^[22,23]^ would not be meaningful due to difference in the employed receptor structure which contains a large part of the N-terminal loop. Our recent analysis of cryo-EM structures demonstrated that, similar to morphine, fentanyl engages the receptor through interactions with residues in SP1 and SP3.^[17]^ We speculate that protonitazene’s interactions within SP2, particularly the *π*-hole bond with Tyr^1.39^, may play a significant role in slowing dissociation relative to fentanyl. A recent cryo-EM study^[14]^ of the *μ*OR-G_i_, *μ*OR-G_z_, and *μ*OR-*β*-arrestin 1 complexes bound to the peptide agonists (DAMGO and endomorphine 1) revealed substantial dynamics of TM1 arising from ligand interactions with SP2 (therein termed TM1-fusion pocket). Importantly, mutations of Tyr^1.39^ differentially modulated *μ*OR signaling bias in a ligand-dependent manner.^[14]^

Our findings suggest that slow dissociation kinetics is a key factor underlying the difficulty of reversing nitazene-related overdoses. Strikingly, the newly published cryo-EM structure^[13]^ of fluetonitazene-bound *μ*OR reproduces the protonitazene–receptor interactions predicted by simulations, including the *π*-hole bond between the nitro group and Tyr^1.39^. The dissociation half-life of fluetonitazene (11.2 min)^[13]^ is shorter than protonitazene, which supports our hypothesis that a bulkier alkoxy group slows dissociation, as fluorine is considerably smaller than a methyl group. Mechanistic insight into how opioid structure governs dissociation kinetics has broad implications, from understanding toxicity to designing more effective countermeasures.

## Supplementary Information

Experimental Materials and Methods. Computational Methods and Protocols. Supplemental Tables (Table S1, S2). Supplemental Figures (Figure S1–S8).

## Data Availability

The data that support the findings of this study are openly available in Github at https://github.com/JanaShenLab/nitazene_kinetics/. Raw trajectories are freely available upon request.

## Acknowledgments

Joseph Clayton is supported by the ORISE fellowship, which is a Research Participation Program at the FDA administered through the Oak Ridge Institute for Science and Education (ORISE) under the agreement between the FDA and Department of Energy. J.S. is supported by the National Institutes of Health (R35GM148261).

## Disclaimer

This article reflects the views of the authors and should not be construed to represent FDA’s views or policies. The mention of commercial products, their sources, or their use in connection with material reported herein is not to be construed as either an actual or implied endorsement of such products by the FDA.

## Entry for the Table of Contents

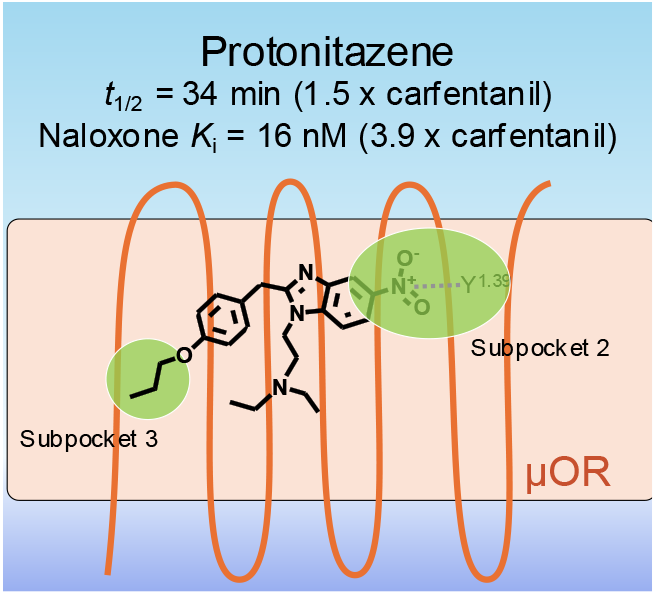

Nitazenes are driving a wave of overdose deaths, but why are they so difficult to reverse? Radioligand experiments point to slow receptor dissociation as a key culprit. In striking agreement with a new cryo-EM structure, simulations reveal specific ligand-receptor interactions, including a unique *π*-hole bond with Y^1.39^, a residue recently shown to modulate *μ*OR signaling bias.

