## Supporting Information for "Slow Dissociation of Nitazenes from the *μ*-Opioid Receptor Underlies the Challenge of Overdose Reversal"

### Contents

**Experimental Materials and Methods** 3

**Computational Methods and Protocols** 5

**Supplemental tables** 9

**Supplemental figures** 10

### Experimental Materials and Methods

*Chemicals.* Protonitazene, etonitazene, and etodesnitazene were purchased from Cayman Chemical Company (Ann Arbor, MI). Other reagents were sourced from Millipore Sigma. The radiolabeling of nitazene compounds was performed by ViTrax, Inc. (Placentia, CA). Specific activities were 21–25, 43 and 38 Ci/mmol, for protonitazene, etonitazene, and etodesnitazene, respectively.

*Cell culture.* Rat C6 glioma cells were transfected with human $\mu$OR cDNA inserted into the plasmid pcDNA3.1neo (cDNA Resource Center, Bloomsburg, PA). Stably transfected cells (C6-h$\mu$OR) were selected with 700 µg/ml G418.

*Membrane preparation.* C6-h$\mu$OR cells were cultured in DMEM supplemented with 10% fetal bovine serum, penicillin/streptomycin, and 400 µg/ml G418. C6-h$\mu$OR cells were grown to confluency, plates were rinsed with calcium-magnesium free phosphate buffered saline (cmf-PBS), and cells were scraped from dishes into cmf-PBS and centrifuged at 1000 x g for 10 min at 4°C. The supernatant was removed. Membranes were washed by the addition of 50 mM Tris buffer (pH 7.4 at 4°C; Tris buffer), homogenized, and centrifuged at 27,000 x g for 20 min at 4°C. This step was repeated. Pellets were covered with 3 ml Tris buffer and frozen at -80°C for later use.

*Saturation binding experiment.* To determine the $K_{\mathrm{d}}$ and $B_{\mathrm{max}}$ values of radioligands at $\mu$OR, saturation binding experiments were conducted with the appropriate incubation times for equilibrium for each radioligand (Figure S1) using six concentrations with duplicate determinations of total and nonspecific binding (10 µM naloxone). The incubation was terminated by rapid filtration through Perkin Elmer Filtermat A filters presoaked in 0.05% polyethylenimine on a Tomtec cell harvester (Hamden, CT) using cold Tris buffer (50 mM, pH 7.4) with a 6-second wash. The filters were dried, spotted with scintillation cocktail, and counted for 2 min after a 4-h interval on a Perkin Elmer microbetaplate counter. Final radioligand concentrations ranged from 0.038–1.99 nM for [$^3$H]protonitazene, 0.003–1.72 nM for [$^3$H]etonitazene, and 0.144–2.80 nM for [$^3$H]etodesnitazene.

*$K_i$ measurement for naloxone and nalmefene.* To determine the $K_{\mathrm{i}}$ of naloxone or nalmefene, competition curves with unlabeled naloxone (0.1 nM–10 µM, half log concentrations) were conducted with each [$^3$H]ligand at concentrations at or above the $K_{\mathrm{d}}$ value for each ligand with the

appropriate incubation time established for each radioligand. The incubation was terminated by rapid filtration as described above.

*Measurement of association and dissociation kinetics.* Association binding assays were performed in 96-well plates using 50 mM Tris buffer (pH 7.4 at 37°C). Nonspecific binding was determined with 10 µM naloxone. Cell membranes (45 to 60 µg protein) were incubated with the radioligand at 37°C at times ranging from 0.08 to 90 min, as appropriate for each radioligand, determined experimentally (Figure S2). The incubation was terminated by rapid filtration as described above.

Dissociation binding assays were performed as described above using the experimentally established equilibration time and the $K_d$ for each radioligand. Triplicate or quadruplicate wells with buffer and cells were incubated at 37°C with [$^3$H]ligand followed by addition of naloxone at times ranging from 0.08 to 90 min, as appropriate for each radioligand, determined experimentally. The incubation was terminated by rapid filtration as described above.

*Data analysis.* GraphPad Prism (version 11.0 for Windows; GraphPad Software, Boston, Massachusetts USA, www.graphpad.com) was used for analyses of receptor saturation and binding assays, ligand-receptor association and dissociation assays.

For receptor saturation binding, data was analyzed for three or more independent assays. A non-linear fit one site analysis of specific bound [$^3$H]ligand (in fmol/mg protein) vs. free [$^3$H]ligand (in nM) was used to determine the equilibrium constant ($K_d$ in nM) and maximal receptor binding sites ($B_{max}$ in fmol/mg protein). The $K_d$ was determined for each independent assay and used to calculate average values. An ordinary one-way ANOVA ($p$=0.0088, $F$=11.52) followed by Tukey's test for multiple comparisons was performed to determine whether nitazene $K_d$ values were significantly different. The $K_d$ for etodesnitazene was significantly higher than protonitazene ($p$ = 0.038) or etonitazene ($p$ = 0.008).

For receptor binding assays to determine $K_i$ (nM) values, the data were normalized to specific binding in the absence of naloxone or nalmefene. Three or more independent assays were measured with duplicate determinations. Data from each independent assay was analyzed using a non-linear sigmoidal dose-response (variable slope) with the 100-to-0 range to determine $IC_{50}$ values and Hill coefficients. $K_\mathrm{i}$ values were calculated using the Cheng-Prusoff transformation: $K_i = IC_{50}/(1 + L/K_\mathrm{d})$, where $L$ is the radioligand concentration and $K_\mathrm{d}$ is the binding affinity of the radioligand, as determined in the saturation binding experiment.

To determine the apparent or observed association rate $k_\mathrm{obs}$ (in $\mathrm{min}^{-1}$), a non-linear one phase exponential analysis was performed on the measured [$^3$H] ligand counts per minute (cpm) from association binding assays (Figure S2). A total of 9 or 10 assays using different [$^3$H]ligand concentrations were generated for each radioligand. Data was collected at time points up to 60 minutes. Linear fits of $k_\mathrm{obs}$ versus radioligand concentration were used to calculate the association rate constant $k_\mathrm{on}$ (in $\mathrm{nM}^{-1}\mathrm{min}^{-1}$) for each radioligand (Figure S3).

Using dissociation assays, the dissociation rate of opioids was characterized by measuring [$^3$H]ligand cpm over time. Data was collected from three or more assays at time points up to 90 min. The half-life of dissociation ($t_{1/2}$) was determined by fitting this data to a one-phase exponential decay model.

### Computational Methods and Protocols

*System preparation.* The structures of $\mu$OR bound to protonitazene, etonitazene, and etodesnitazene were taken from our previous modeling work.[1] Briefly, using three docking programs (Glide,[2] MOE[3] and AutoDock[4]), a series of nitro-containing and nitro-less nitazenes (including the three studied here) were docked to an active conformation of $\mu$OR using the co-crystal structure of $\mu$OR:BU72 complex (PDB 5C1M)[5] or the cryo-EM structure of $\mu$OR:mitragynine pseudoindoxyl complex (PDB 7T2G[6]) as a template. The docking resulted in three types of binding poses, based on whether the nitazene's benzimidazole is placed in the receptor's subpocket 1 (SP1, between transmembrane helices TM2 and TM3), subpocket 2 (SP2, between TM1, TM2, and TM7), or subpocket 3 (SP3, between TM5 and TM6). The stabilities of these poses were then analyzed using binding-pose metadynamics, conventional MD (cMD), and metadynamics-based free energy simulations, which led to the conclusion that the SP2 binding pose is most stable.

The simulation system was prepared using CHARMM-GUI.[7] The $\mu$OR-nitazene complex structure was oriented in a bilayer of POPC lipids (roughly 175 lipids per leaflet) using PPM 2.0 web server.[8] Water molecules were added with a layer thickness of 22.5 Å. 78 sodium and 92 chloride ions were added to neutralize the system at pH 7.4 (assuming default protonation states for acidic and basic residues) and achieve an ionic strength of 150 mM. All histidines were set to the HID tautomer, including H297$^{6.52}$ in the orthosteric binding site which was shown to affect fentanyl's residence time at the receptor.[9] The protein and lipids were represented by the CHARMM36m protein[10] and CHARMM36 lipid force fields,[11] respectively, while water was represented by the CHARMM modified TIP3P model.[12,13] The force field parameters of sodium and chloride ions were taken from Refs[14,15] with the atom-pair specific adjustment (also known as NBFIX) to reduce overbinding of sodium to carboxylates.[16] The CHARMM modified TIP3P model was used to represent water.[12,13] The force field parameters of nitazenes were assigned using CHARMM-GUI,[7] which utilizes the CGenFF algorithm for assigning force field parameters based on analogy to compounds with optimized parameters.[17,18]

Our previous work followed the default preparation protocol of CHARMM-GUI,[7] which involved minimizing over 5,000 steps, followed by heating under constant volume to 300 K over 125 ps with restraints on the protein positions, lipid positions, and lipid dihedral angles. Restraints were gradually reduced over 2.25 ns under constant pressure, then 100 ns was conducted to refine the docked position of the nitazene. In this work, we further refined the docked position by conducting an additional 200 ns for $\mu$OR in complex with protonitazene, etonitazene, or etodesnitazene. Analysis of trajectories was conducted using CPPTRAJ.[19]

*Metadynamics simulations.* Following 300-ns MD simulation, 15 independent well-tempered metadynamics[20] simulations were conducted. Following our previous work,[21] two collective variables (CV) were used: the $z$-position of the center of mass (COM) of the nitazene relative to that of the orthosteric binding pocket, and the coordination number (CN). The COM of the binding pocket was defined as using the $C_\alpha$ atoms of residues Y75, Q124, N127, W133, L144, D147, Y148,

M151, F152, L232, K233, V236, A240, W293, I296, H297, V300, W318, H319, I322, and Y326. The CN represents the number of contacts between the nitazene and $\mu$OR, defined as

$$CN = \sum_{i \in \mu\text{OR}} \sum_{j \in \text{nitazene}} \frac{1 - (d_{ij}/d_{\text{cut}})^8}{1 - (d_{ij}/d_{\text{cut}})^{16}},$$

where $d_{ij}$ is the distance between the heavy atoms $i$ in $\mu$OR and $j$ in the nitazene, and $d_{cut}$ is the cutoff distance (set to 4.5 Å). Following our previous dissociation simulations,[21] each simulation was conducted until an unbound state was reached, defined as the z-position CV reaching 15 Å. The Gaussian weight parameter was set to 0.5 kcal/mol and the width was set to 0.5 Å for the z-position CV and 5 for the coordination CV. Hills were deposited every 20 ps, while the tempering parameter for well-tempered metadynamics was set to 14.

OpenMM v8.0[22] was used to run both the preparation protocol and the well-tempered metadynamics simulations; PLUMED v2.7[23] was used to control the metadynamics, via the OpenMM PLUMED plugin. Pressure was maintained at 1 bar using the Monte-Carlo barostat[24] with a coupling frequency of 100 steps, and with the x and y dimensions coupled isotropically. Temperature was controled by the Langavin thermostat[25] with a friction coefficient of 1. SETTLE[26] was used for water molecules, while SHAKE[27] was used for all other atoms.

*Estimation of residence times at the receptor.* Infrequent metadynamics[28,29] was used to estimate the nitazene-$\mu$OR residence time. For each metadynamics simulation, the real exit time $t_{exit}$ was estimated by multiplying the simulation time (accelerated by metadynamics) by the running temporal average of the metadynamics potential, called the acceleration factor $\alpha$[28]

$$\alpha =< \exp(\beta V_{meta}(t)) >$$

where $V_{meta}(t)$ is the bias potential at simulation time t. Following previous work[21,30] the $t_{exit}$ of each fifteen metadynamics simulations was then used to form an empirical cumulative distribution function (CDF) and fit this to a theoretical CDF for a Poisson process, representing the probability of observing at least one dissociation event by time $t$:

$$P_{n\geq 1}(t) = 1 - \exp(-t/\tau)$$

where $\tau$ is the residence time. To estimate the errors of $\tau$ we performed bootstrapping analysis where the fifteen $t_{exit}$ values were resampled 10,000 times. Each resampling was used to fit a Poisson process, and the Kolmogorov-Smirnov (KS) test was used to test if the sample and fitted Poisson process share the same underlying distribution.[30] If the sample showed a p-value lower than 0.05, then the sample was discarded. Remaining samples (where the p-value was equal or greater than 0.05) were used to calculate the mean and standard error of $\tau$.

To produce unbiased probability distributions and consequentially the potential of mean force, frames were reweighted through an in-house script that follows the REWEIGHT_BIAS module implemented in PLUMED.[23]

### Supplemental tables

Table S1: Experimentally determined $t_{1/2}$ (min) of nitazenes and fentanyls at different concentrations of unlabeled naloxone

| Naloxone | 10 nM | 100 nM | 1 $\mu$M | 10 $\mu$M |
|---|---|---|---|---|
| Protonitazene | 34.46±2.68 | 25.87±1.12 | 27.89±3.38 | 23.52±1.90 |
| Etonitazene | 14.15±0.64 | 10.87±0.14 | 10.30±0.10 | |
| Etodesnitazene | 8.25±1.13 | 6.34±0.18 | 5.37±0.50 | |
| Fentanyl | 4.27±0.53 | | | 3.67±0.48 |
| Carfentanil | 22.9±1.8 | | | 19.72±1.64 |

Three replicate experiments were conducted for each compound and naloxone concentration. Average and standard error are shown. For fentanyl and carfentanil, two naloxone concentrations were used in experiments.

Table S2: Simulation estimated $t_{1/2}$ for nitazenes. Each receptor-ligand system was subject to 15 independent metadynamics simulations, yielding 15 estimates of the unbiased dissociation times. The distribution was then fit to a theoretical Poisson process to obtain $t_{1/2}$. For estimation of statistical uncertainty, 10,000 bootstrapping trials were performed, keeping only trials that yielded a $p$-value above 0.15 (valid trials).

| Compound | $t_{1/2}$ (s) | # Valid trials | Avg. p value |
|---|---|---|---|
| Protonitazene | 18.3±0.3 | 6293 | 0.15 |
| Etonitazene | 0.26±0.02 | 7042 | 0.15 |
| Etodesnitazene | 0.0144±0.0006 | 4472 | 0.11 |
| Fentanyl | 1.469±0.017 | 6972 | 0.18 |

#### Supplemental figures

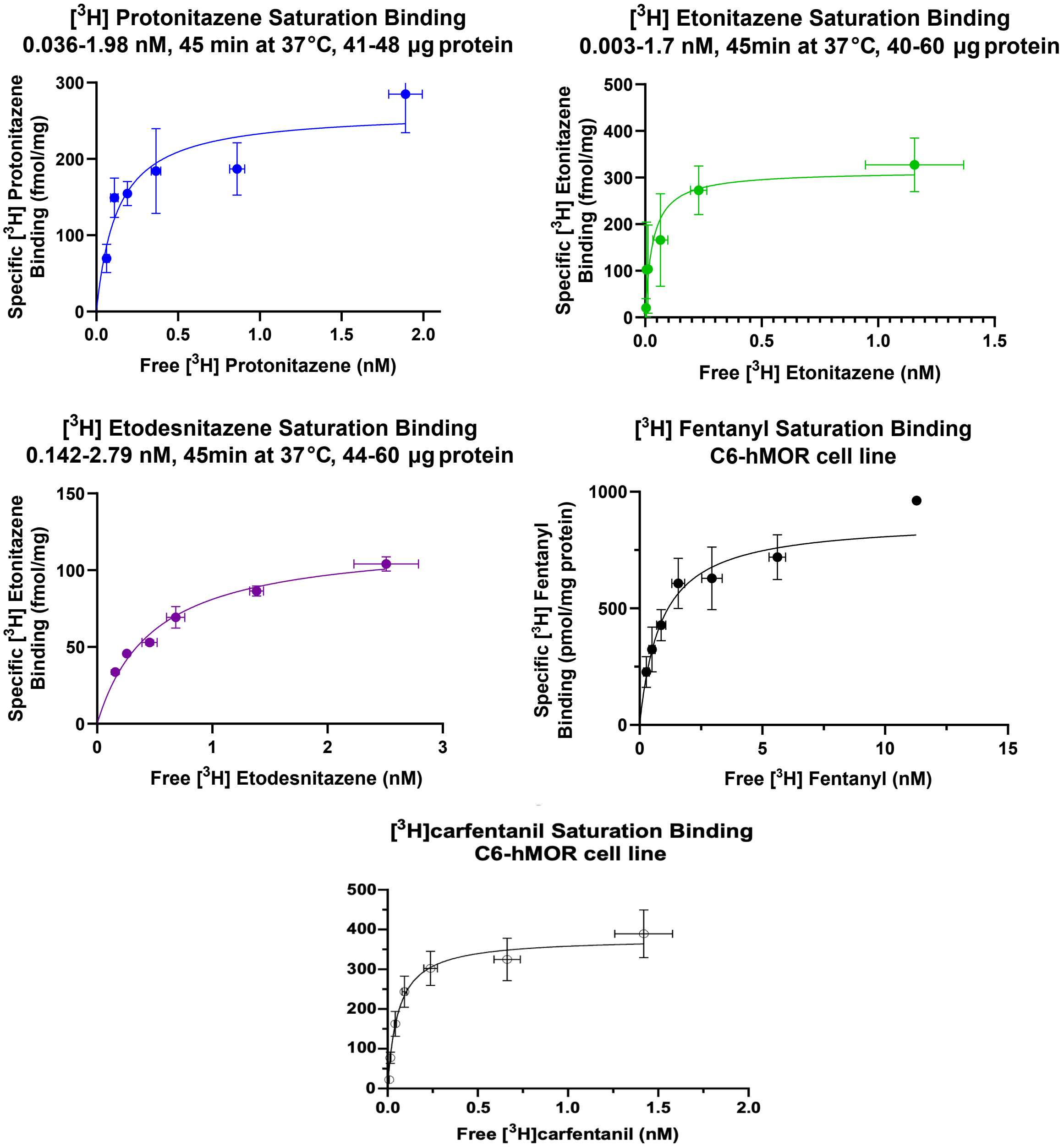

Figure S1: **Composite saturation binding curves for [$^3$H] nitazenes and [$^3$H] fentanyl binding to $\mu$OR**. Composite curves were generated using average specific bound [$^3$H]ligand vs. average free [$^3$H]ligand (nM) from 3 or 5 independent assays. Curves are the best fits to the one site specific binding model. Bars represent the standard error of the mean.

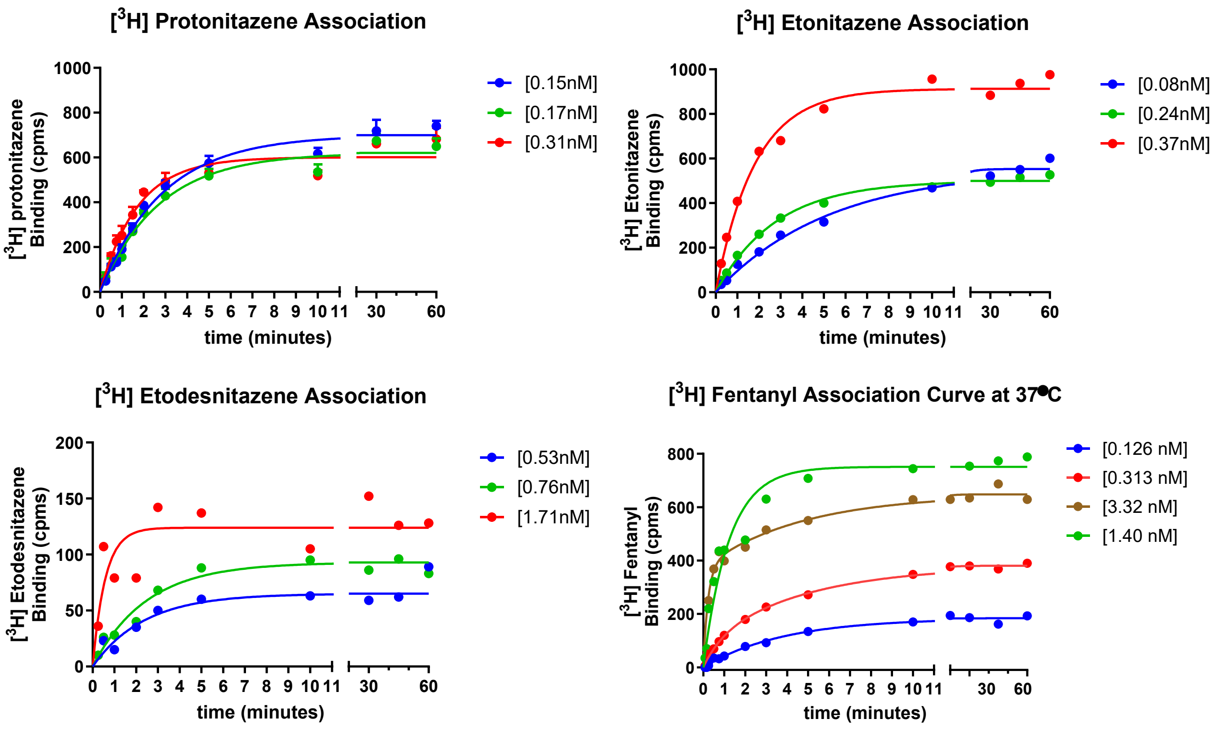

Figure S2: **Representative association curves showing the time course for association of [$^3$H]radioligands to the $\mu$OR**. One phase exponential (pseudo-first order) association curves corresponding to each association assay are shown. A total of 3 or 4 curves using different [$^3$H] radioligand concentrations are shown.

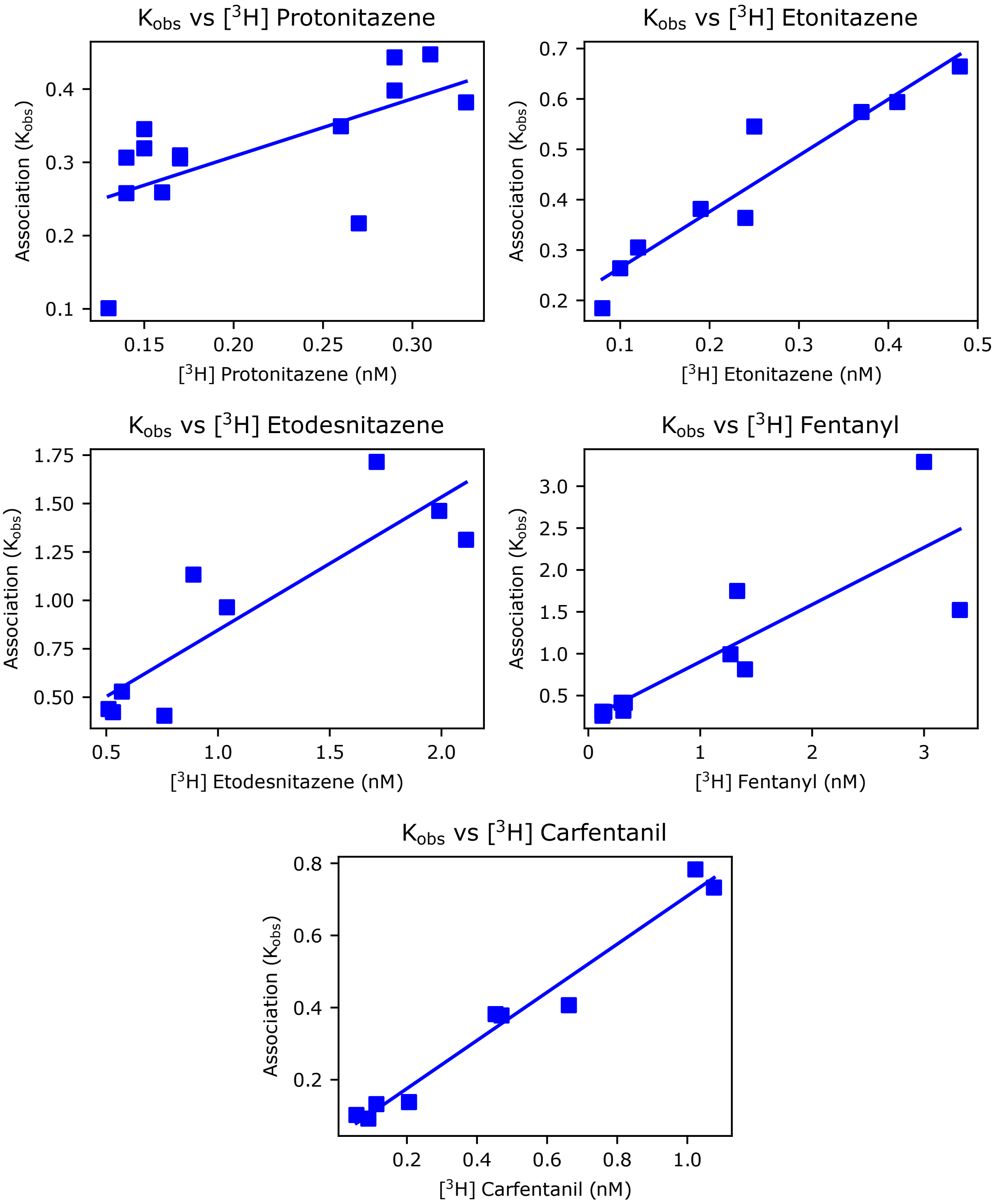

Figure S3: **Apparent association rate for [$^3$H]nitazenes and [$^3$H]fentanyl**. For each association assay, a one phase exponential association curve (see Figure S2) was used to calculate the apparent or observed association kinetic rate ($k_{obs}$ in min$^{-1}$) at each radioligand concentration. A total of 9 or 13 concentrations were used. The panel above shows $k_{obs}$ plotted versus the concentration of unbound radioligand (nM). Linear regression of the data yields the estimated $k_{on}$ in nM$^{-1}$min$^{-1}$ (slope of the line) for each radioligand.

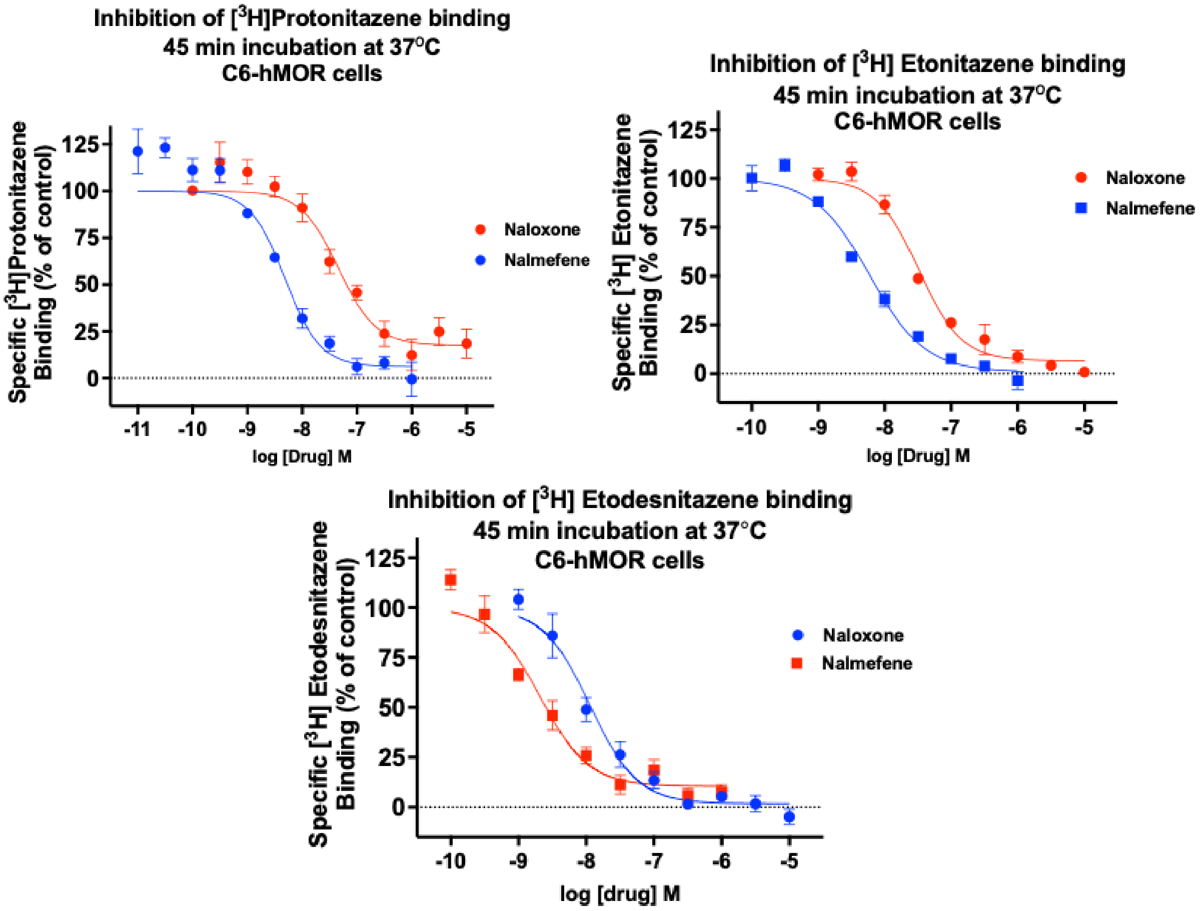

Figure S4: Competitive displacement of a nitazene due to naloxone or nalmefene at the $\mu$OR.

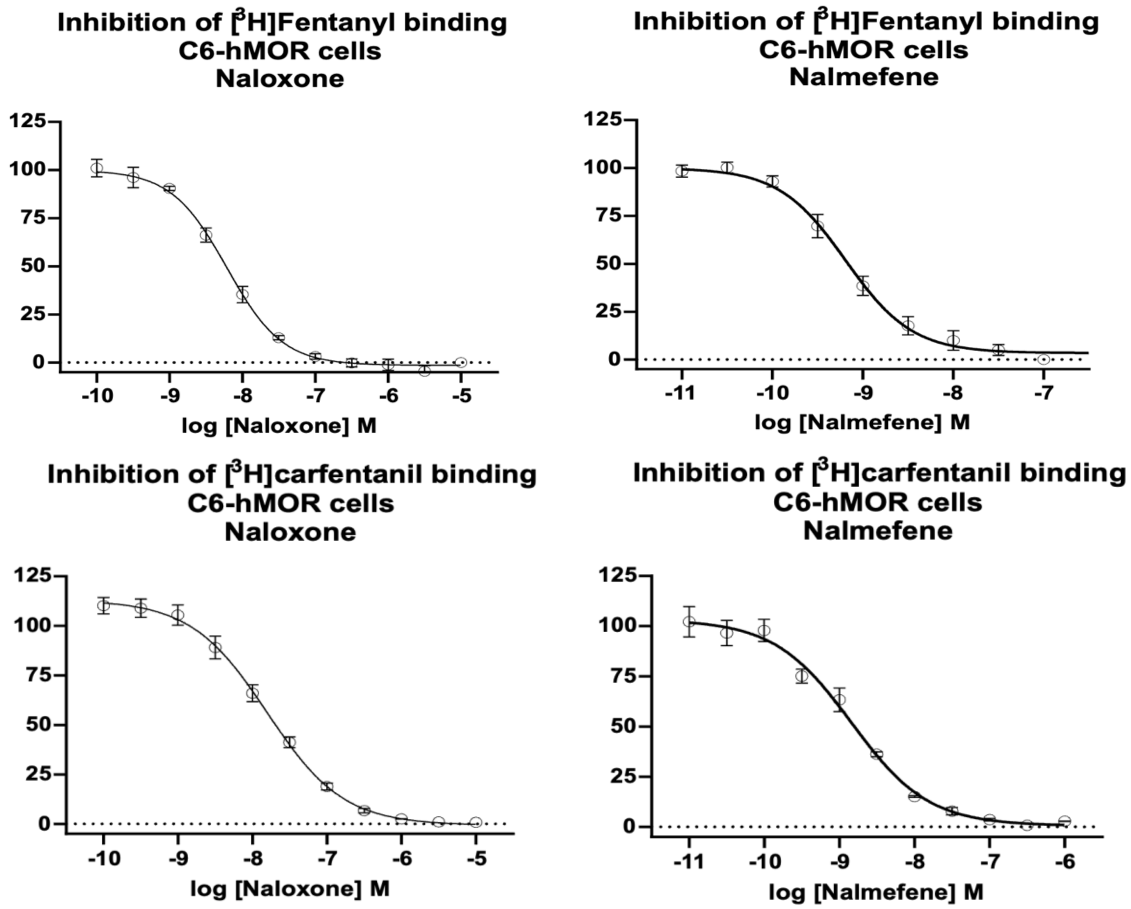

Figure S5: Competitive displacement of either fentanyl or carfentanyl due to naloxone or nalmefene at the $\mu$OR.

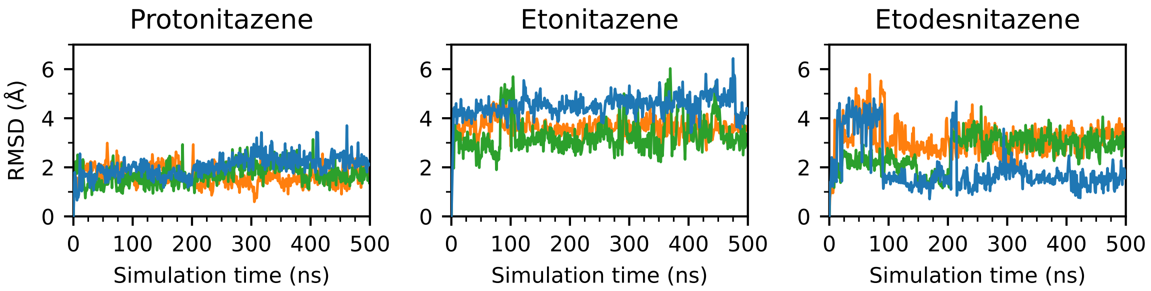

Figure S6: Root mean squared deviation (RMSD) of heavy atoms of the ligand during the three 500 ns conventional MD simulation, with respect to the docked structure. The starting conformation for estimating the dissociation $t_{1/2}$ was taken at 300 ns of the simulation show in blue.

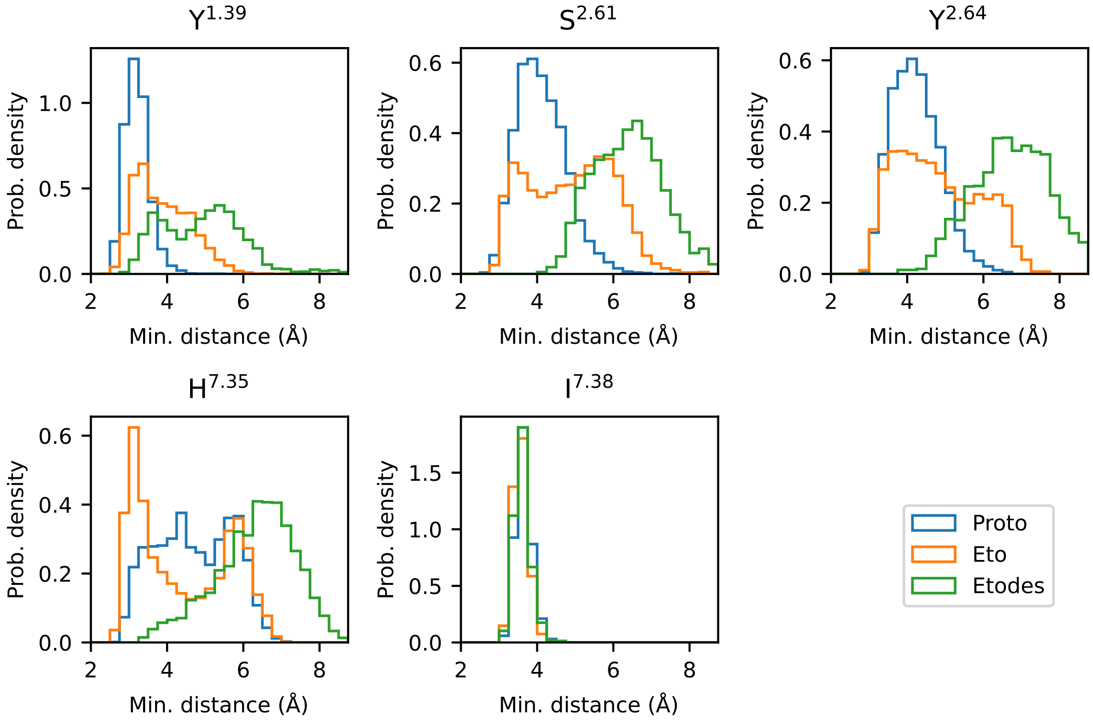

Figure S7: Minimum distance between non-hydrogen atoms of each nitazene and residues within SP2. Only the final 100 ns of three 300 ns simulations were analyzed.

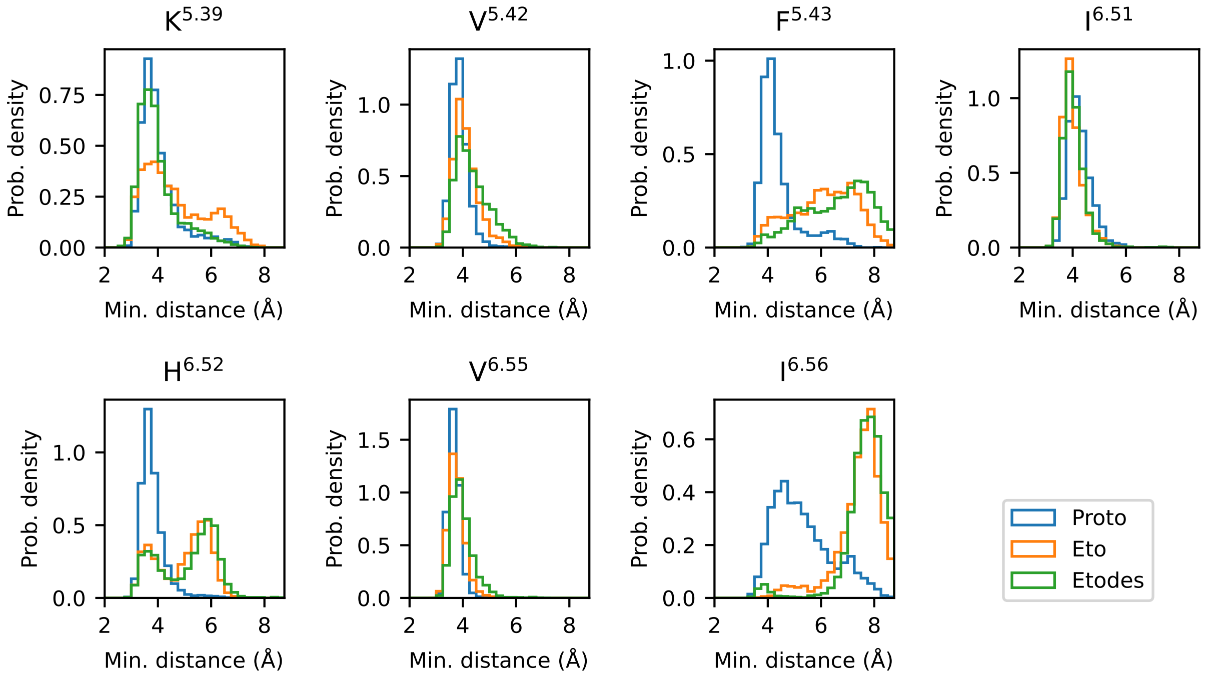

Figure S8: Minimum distance between non-hydrogen atoms of each nitazene and residues within SP3. Only the final 100 ns of three 300 ns simulations were analyzed.
